# Endothelial Lipase Variant, T111I, Does Not Alter Inhibition by Angiopoietin-like Proteins

**DOI:** 10.1101/2023.08.18.553740

**Authors:** Kelli L. Sylvers-Davie, Kaleb C. Bierstedt, Michael J. Schnieders, Brandon S. J. Davies

## Abstract

High levels of HDL-C are correlated with a decreased risk of cardiovascular disease. HDL-C levels are modulated in part by the secreted phospholipase, endothelial lipase (EL), which hydrolyzes the phospholipids of HDL and decreases circulating HDL-C concentrations. A 584C/T polymorphism in *LIPG*, the gene which encodes EL, was first identified in individuals with increased HDL levels. This polymorphism results in a T111I point mutation the EL protein. The association between this variant, HDL levels, and the risk of coronary artery disease (CAD) in humans has been extensively studied, but the findings have been inconsistent. In this study, we took a biochemical approach, investigating how the T111I variant affected EL activity, structure, and stability. Moreover, we tested whether the T111I variant altered the inhibition of phospholipase activity by angiopoietin-like 3 (ANGPTL3) and angiopoietin-like 4 (ANGPTL4), two known EL inhibitors. We found that neither the stability nor enzymatic activity of EL was altered by the T111I variant. Moreover, we found no difference between wild-type and T111I EL in their ability to be inhibited by ANGPTL proteins. These data suggest that any effect this variant may have on HDL-C levels or cardiovascular disease are not mediated through alterations in these functions.

## INTRODUCTION

High density lipoprotein plays an important role in cholesterol homeostasis by participating in reverse cholesterol transport, a process in which excess cholesterol is loaded onto HDL to be removed from peripheral tissues and macrophages then transported back to the liver for excretion (1). Reverse cholesterol transport, in combination with additional anti-atherogenic properties of HDL, including anti-inflammatory, anti-apoptotic, and anti-thrombotic properties, are like responsible for the correlation between HDL levels and cardiovascular health (2–5). Endothelial lipase (EL), a secreted lipase derived from endothelial cells in highly vascularized tissues, hydrolyzes the phospholipids that make up the outer shell of HDL at the sn-1 position (6). This remodeling of the HDL membrane accelerates clearance of HDL by downstream receptors, decreasing plasma HDL-cholesterol (HDL-C) levels (7). The link between EL and HDL levels has been demonstrated and validated in mice, where EL-deficiency results in increased HDL-C and HDL phospholipids, and overexpression of EL results in decreased HDLC (8–10).

Several polymorphisms in *LIPG*, the gene that encodes endothelial lipase, have been identified in human patients. A common polymorphism located in exon 3, 584C/T (rs2000813), results in an amino acid substitution, Thr111Ile (11). The T111 residue is conserved in humans, primates, and rats, but not in mice or rabbits. Although this polymorphism was first identified in a 2002 study of individuals with elevated HDL-C levels (11), subsequent studies have produced mixed results when attempting to link this variant to altered HDL concentrations. Several studies have found an association between Thr111Ile and increased HDL levels and reduced risk of coronary heart disease risk (10, 12–17), but other studies failed to find any correlation (18–23). In biochemical studies, EL T111I appears to have similar phospholipase activity as wild-type EL (23, 24). Thus, if and how this mutation affects HDL levels and other cardiac risk factors remains unclear.

The major endogenous regulator of EL activity is the hepatokine, angiopoietin-like 3 (ANGPTL3). By itself, ANGPTL3 is a potent inhibitor of EL (25), and when in complex with ANGPTL8, can also potently inhibit lipoprotein lipase (LPL), a closely related lipase to EL, (26–31). ANGPTL3 inhibition of EL results in increased levels of HDL and HDL-C (25, 32). An altered ability of EL T111I to be inhibited by ANGPTL3 could explain the differences in HDL-C levels sometimes observed with this allele. However, the ability of ANGPTL3 to inhibit the activity of the T111I variant of EL has not yet been examined.

In this study, we evaluated the EL variant with a T111I substitution for phospholipase activity, stability, and ability to be inhibited by ANGPTL proteins.

## MATERIALS AND METHODS

### Expression constructs

The generation of plasmid constructs expressing strep-tagged mouse ANGPTL3 (pHS18) and strep-tagged human ANGPTL3 (pHS15) have been described previously (26). The generation of the plasmid construct expression V5-tagged human ANGPTL4 (pHS2) has been described previously (33).

A plasmid expressing untagged human EL (pKS6) was generated by amplifying the coding sequence of human EL from cDNA (LIPG; MGC MHS6278-202806078) and inserting it into the vector pCDNA6 using InFusion cloning (Clontech). An EL containing a T111I mutation (pKS17) was generated by introducing a *332C > T* substitution into the EL coding region of pKS6 using site-directed mutagenesis. A lentiviral construct expressing EL with a T111I mutation (pKS18) was generated by introducing a *332C > T* substitution into the EL coding region of pKS7, a lentiviral construct expressing wild-type EL (34). Lentiviruses with this construct were produced by transfecting human embryonic kidney 293T (HEK 293T) cells with pKS18 and the lentiviral packaging vectors pMD2.G (Addgene plasmid #12259; http://n2t.net/addgene:12259; RRID:Addgene 12259), pRSV-Rev (Addgene plasmid #12253; http://n2t.net/addgene:12253; RRID:Addgene 12253)(35), and pMDLg/pRRE (Addgene plasmid #12251; http://n2t.net/addgene:12251; RRID:Addgene 12251)(35). All lentiviral packaging vectors were gifts from Didier Trono. Lentivirus-containing media was collected, concentrated using Lenti-X Concentrator (Clontech; catalog no. 631231), and used to generate EL-expressing cell lines as described below. A construct expressing full-length V5 tagged mouse ANGPTL4 (pHS5) was generated by amplifying full-length mouse ANGPTL4 cDNA (OpenBiosystems) and inserting it into a pCDNA6 vector using In-Fusion cloning (Clontech). A V5 tag was then appended to the C terminus of the open reading frame using Phusion site-directed mutagenesis (New England Biolabs).

### Cell lines

HEK 293T cells were grown and maintained in DMEM supplemented with 5% FBS, 1% Penicillin/streptomycin antibiotics, and 1% L-Glutamine (complete DMEM).

Rat heart microvessel endothelial cells (RHMVECs; VEC Technologies, Inc) were grown in MCDB-131 base medium (GenDEPOT) supplemented with 10 mM L-glutamine, 1% penicillin/streptomycin antibiotic solution (10,000 U/ml penicillin and 10,000 μg/ml streptomycin; Gibco), 5% fetal bovine serum (Atlanta Biologicals), 1 μg/ml hydrocortisone (Sigma-Aldrich), 10 μg/ml human epidermal growth factor (Gibco, Life Technologies), and 12 μg/ml bovine brain extract (Lonza). To generate a stable cell line expressing human EL (T111I), 80% confluent RHMVECs in 6-well plates were transduced with human T111I EL (pKS18) lentiviruses with 4 μg/ml Polybrene (#134220; Santa Cruz Biotechnology) in a total volume of 1 ml DMEM. Twenty-four hours post-transduction, cells were washed with PBS and incubated in MCDB-131 complete medium for 48 h. Cells were then subjected to selection with 6 μg/ml puromycin for 5 days. RHMVECs stably expressing wild-type human EL (pKS7RHMVECs) were generated as described previously (34).

### Production and quantification of ANGPTL3 and ANGPTL4 conditioned media

To produce human and mouse ANGPTL3 and ANGPTL4 protein, HEK 293T cells were grown to 80% confluency in T25 flasks in complete DMEM. Cells were transfected with 5 μg of pHS18 (strep-tagged mouse ANGPTL3), pHS15 (strep-tagged human ANGPTL3), pHS5 (V5tagged mouse ANGPTL4), or pHS2 (V5-tagged human ANGPTL4) and 10 μl of 1 mg/ml PEI (polyethylenimine). Cells were switched to serum-free DMEM and 1X ProteaseArrest protease inhibitor cocktail (APExBIO) 24 h post-transfection. Conditioned media were collected 48–72 h later. To produce 293T control media (CM), HEK293T cells were transfected and media collected in the same manner, only no DNA was added to the transfection. This control conditioned media was used as a control for all experiments involving ANGPTL conditioned media. The concentration of ANGPTL3 in conditioned media was determined via western quantification against a batch of mouse ANGPTL3 with a known concentration, previously quantified via ELISA kit (RayBiotech, catalog #ELM-ANGPTL3-1). The concentration of ANGPTL4 in conditioned media was determined via western quantification against a batch of human ANGPTL4 with a known concentration, previously quantified via a human ANGPTL4 ELISA kit (Sigma).

### Production and quantification of endothelial lipase conditioned media

To produce EL conditioned media, HEK 293T were grown to 80% confluency in DMEM supplemented with 5% FBS, Penicillin/streptomycin antibiotics, and L-Glutamine. Cells were transfected with 10 μg of pKS6 (human WT EL) or pKS17 (human T111I EL) and 20 μl of 1 mg/ml PEI (polyethylenimine). Media was changed to serum-free OptiMEM containing 1X ProteaseArrest and 0.1 U/ml heparin (Fresenius Kabi USA, LLC) 24 hours post-transfection. EL conditioned media was collected after an additional 24 hours. Protein expression was confirmed by western blotting and tested via phospholipase activity assay. EL concentration was determined by comparing EL protein signal with a quantified batch of strep-tagged human EL protein via quantitative western blot.

### Western blot

Protein samples were size fractionated on 12% SDS-PAGE gels and then transferred to a nitrocellulose membrane. Membranes were blocked with casein buffer (1% casein, Fisher Science Education). Primary antibodies were diluted in casein buffer + 0.1% Tween. Primary antibody dilutions were 1:3000 for a mouse monoclonal antibody against EL (LIPG antibody clone 3C7, Lifespan Biosciences), 1:2000 for a goat antibody against beta-actin (Abcam), 1:3000 for a rabbit polyclonal antibody against Strep-tag II (Abcam) to detect Strep-tagged ANGPTL3, or 1:6000 for a mouse monoclonal antibody against V5 tag (R960-25; Invitrogen) to detect V5tagged ANGPTL4. After washing with PBS-T, membranes were incubated with Dylight680- or Dylight800-labeled secondary antibodies (Thermo Scientific) diluted 1:5000 in casein. After washing with PBS-T, antibody binding was detected using an Odyssey Clx Infrared Scanner (Li-Cor).

### EL activity assays

Phospholipase activity was monitored using the EnzChek Phospholipase A1 assay kit (ThermoFisher) as described previously (36). EL and ANGPTL3 or ANGPTL4 proteins in conditioned media were combined and incubated at 37°C for 30 minutes. Following incubation, 50 μl of sample was mixed with 50 μl of substrate solution and were incubated at room temperature (approximately 20-22°C) for 30 minutes, reading fluorescence (485 nm excitation/515 nm emission) every 1 or 2 minutes with an Infinite F200 plate reader (Tecan). Relative phospholipase activity was calculated by calculating the slope of the linear part of the curve (typically in the range between 5 to 25 minutes) and then subtracting out the slope of the blank (sample with no EL conditioned media).

To detect phospholipase activity on the cell surface, RHMVECs expressing WT EL (pKS7-RHMVEC) or T111I EL (pKS18-RHMVEC) were grown to confluency in 96-well clear bottom plates coated with fibronectin. After washing twice with 1× sterile PBS, cells were incubated with 50 μl of ANGPTL3, ANGPTL4, or control conditioned media for 30 min at 37°C. After incubation, 50 μl of EnzChek Phospholipase A1 assay substrate was added, and fluorescence (485 nm excitation/515 nm emission) was read at room temperature (approximately 20–22°C) every 2 min for 30 min with an Infinite F200 plate reader. Relative phospholipase activity was determined by calculating the slope of the linear part of the curve (typically in the range between 5 and 25 min) and then subtracting out the slope of samples containing untransduced RHMVECs incubated with 50 μl control conditioned media. Each condition was normalized as a percentage of their respective positive control (either WT or T111I EL expressing cells which did not receive any ANGPTL conditioned media).

### EL activity assay with HDL substrate

EL conditioned media and 67 or 116.4 mg/dL (cholesterol) of human HDL (Lee Biosolutions) were incubated together for 60 minutes at 37°C. After incubation, 50 μL of sample was loaded into a Greiner clear 96-well plate. NEFA levels in each sample were measured using a commercial kit (HR series NEFA-HR, Wako) and following the manufacturer’s instructions. Briefly, 112.5 μL of NEFA Reagent A (0.53 U/mL Acyl-coenzyme A synthetase, 0.31 mmol Coenzyme A, 4.3 mmol/L Adenosine triphosphate, 1.5 mmol/L 4-aminoanitipyrine, 2.6 U/mL Ascorbate oxidase, 0.062% Sodium azide) was added to each sample and incubated for 10 minutes at 37°C. 37.5 μL of NEFA Reagent B (12 U/mL Acyl-coenzyme A oxidase, 14 U/mL Perosidase) was then added and samples were incubated for 5 minutes at 37°C. Absorbance was read at 560 and 670 nm with a Spectramax i3 (Molecular Devices). For data analysis, the values read at 670 nm were subtracted from 560 nm values.

### Predicted structure modeling

The protein structure was predicted using ColabFold (37), an adaptation of AlphaFold2 (38) which uses a slightly different algorithm to generate the multiple sequence alignment. The EL FASTA sequence was acquired from the Uniprot database entry Q9Y5X9 (39). This sequence was used as an input to generate the MSA used by the AlphaFold2 algorithm. Five predicted structures were generated from the algorithm and the structure with the highest confidence rank was selected to be the representative structure. Figures were generated using Pymol by importing the pdb files generated from ColabFold.

### Statistics

Statistics were performed using GraphPad Prism 9. Single point lipase activity assays comparing EL and EL T111I were analyzed using unpaired student t tests. All dose-response and time-course activity curves were compared using one phase decay least squares fit.

## RESULTS

### Phospholipase activity of EL T111I

There are conflicting reports of the association of the EL T111I variant with HDL levels (10–20, 22, 23). We reasoned that if the T111I mutation does alter HDL-C levels, these alterations could be mediated by decreases in EL activity or protein stability, or an increase in susceptibility to ANGPTL3 inhibition.

To address the potential effect of the T111I mutation on EL function, we generated expression constructs coding for wild-type and T111I mutant EL **(Fig. 1A)**. When these constructs were transiently expressed in HEK 293T cells, we found that expression, secretion, and cleavage of the EL T111I mutant was similar to wild-type EL **(Fig. 1B)**. Consistent with a previous report (23), we also found that phospholipase activity of the T111I mutant was similar to wild-type, whether using an artificial substrate **(Fig. 1C)**, or when measuring hydrolysis of phospholipids of human HDL (**Fig. 1D**).

**Figure 1.**
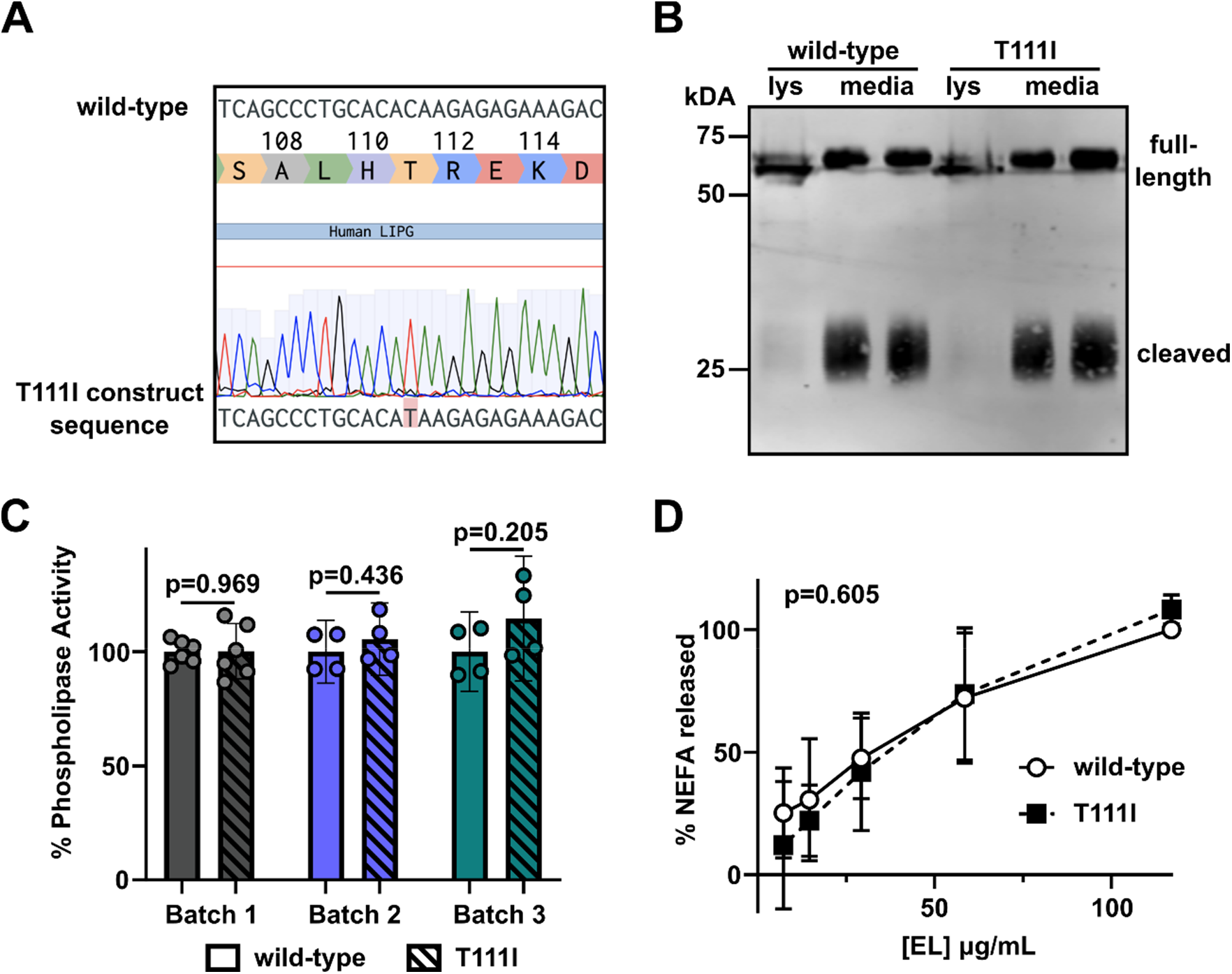
Expression and activity of the EL T111I mutation. **(A)** Sequencing data showing successful generation of the 584C/T mutation in human LIPG. **(B)** Expression and secretion of wild-type and T111I EL. Western blot shows expression of EL in the lysate (lys) and media of 293T cells transfected with wild-type or T111I EL. **(C)** Activity of three independently collected batches of WT and T111I EL. For each batch constructs expressing WT and T111I EL were transfected at the same time. After matching protein levels via western blot intensity, phospholipase activity of the conditioned media was measured using a fluorescence-based phospholipase activity assay. Bars show mean ±SD. P values determined using student t test. **(D)** Phospholipase activity of EL as measured by the release of NEFAs from HDL. The indicated concentrations of EL were combined with human HDL for 60 min at 37°C. Points represents 4 independent experiments using two different batches of human HDL (mean ±SD). P value determined by one phase decay least squares fit.

### Stability and structure of EL T111I

The structure of EL has not been solved. We utilized ColabFold (37), a derivative of Alphafold2 (38), to predict the structure of wild-type human EL **(Fig. 2A)**. In agreement with previously published predictions (40), the T111 residue is located at the end of an alpha helix in the N-terminus domain of EL. In tandem with the predicted structure, we utilized Dynamut2 (41) to predict how the T111I mutation might alter the stability of EL. The predicted ΔΔG^Stability^ was −0.18 kcal/mol, a value indicative of a negligible change. We have previously observed that wild-type EL spontaneously loses activity at 37°C (34), a trait shared with the closely related lipase, lipoprotein lipase (42, 43). Therefore, we asked if the T111I EL spontaneously lost activity at a different rate than wild-type LPL. However, we found that, at 37°C, EL with the T111I mutation lost activity at a similar rate compared to wild-type **(Fig. 2B)**.

**Figure 2.**
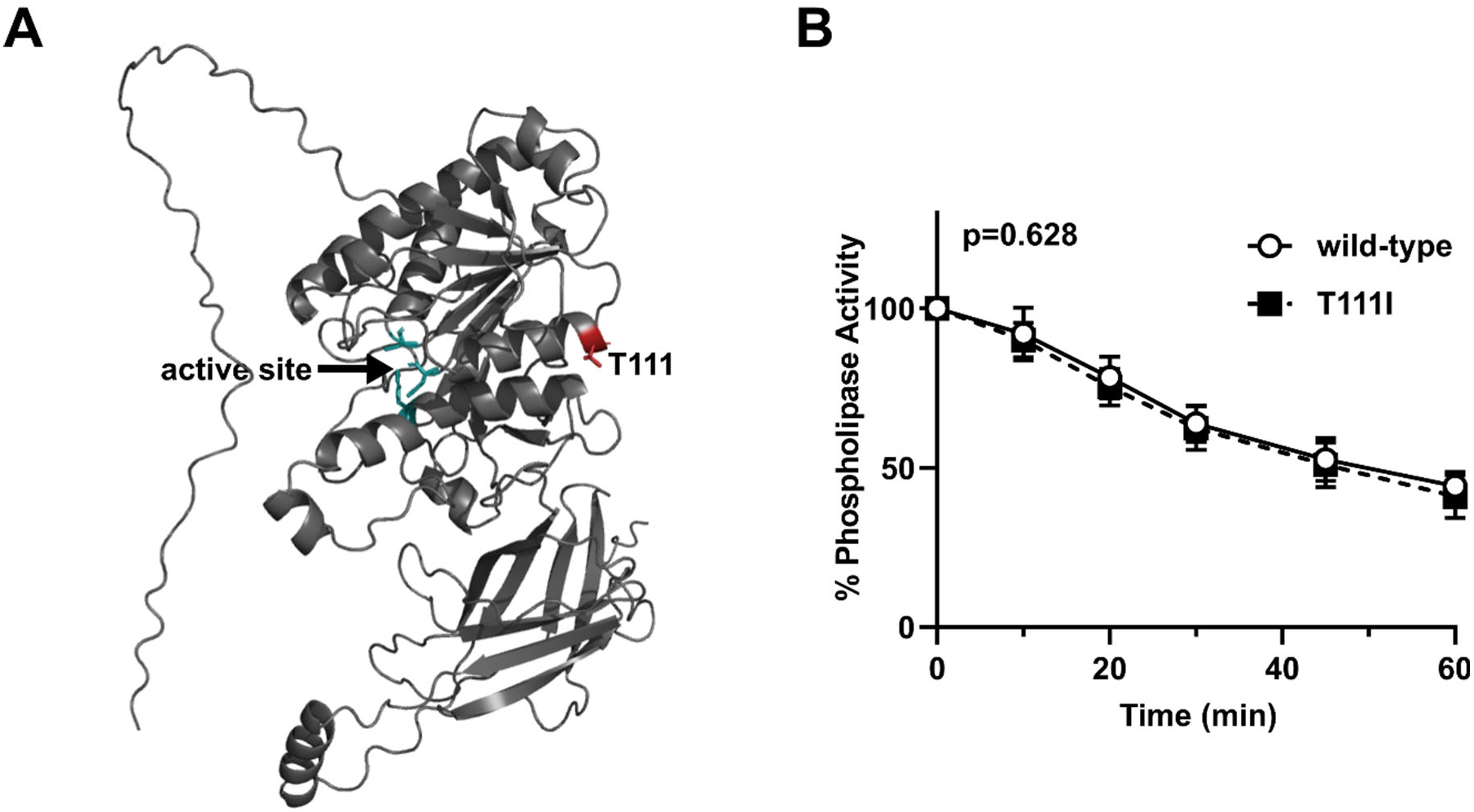
Stability of EL T111I. **(A)** Structure of WT EL as predicted by ColabFold. The catalytic triad of EL is depicted in green and the residue T111 is colored in red. **(B)** Activity over time of WT and T111I EL conditioned media incubated at 37°C. Activity at each time point was measured using a fluorescence-based phospholipase activity assay. Points represents 5 independent experiments (mean ±SD). P value determined by one phase decay least squares fit.

### Inhibition of EL T111I by angiopoietin-like proteins

EL shares 44% homology with the closely related lipase, lipoprotein lipase (LPL). LPL is inhibited by both ANGPTL3, in the form of ANGPTL3/8 complexes, and by ANGPTL4 (26–28, 43–51). The interaction between LPL and ANGPTL4 has been extensively studied and potential binding sites between the two proteins has been mapped through deuterium exchange studies (42, 52).

Because EL is closely related to LPL, we asked if the T111 mapped to an area homologous an ANGPTL4 binding domain in LPL. We aligned the protein sequence of human EL against human LPL using the Clustal format alignment by MAFFT (53) **(Fig. 3A)**. When comparing homology between EL and LPL, the T111 residue aligns with one of the binding site clusters to which ANGPTL4 binds (**Fig. 3A,B**). Therefore, it is possible that ANGPTL4 and/or ANGPTL3 interacts with EL at the same site, and that the T111I mutation could disrupt this interaction. To test this idea, we first measured EL phospholipase activity after incubation with increasing concentrations of human or mouse ANGPTL3 protein. Inhibition of both wildtype and T111I EL was dose-dependent, consistent with our previous findings (34). Moreover, both human and mouse ANGPTL3 inhibited T111I EL to a similar level as wild-type EL **(Fig. 4A-B)**. Recently ANGPTL4 has also been reported to inhibit EL (54). Therefore, we asked if ANGPTL4 inhibition was affected by the T111I mutation. As with ANGPTL3, there was little difference in inhibition of T111I EL by ANGPTL4 when compared to wild-type EL **(Fig. 4C-D)**.

**Figure 3.**
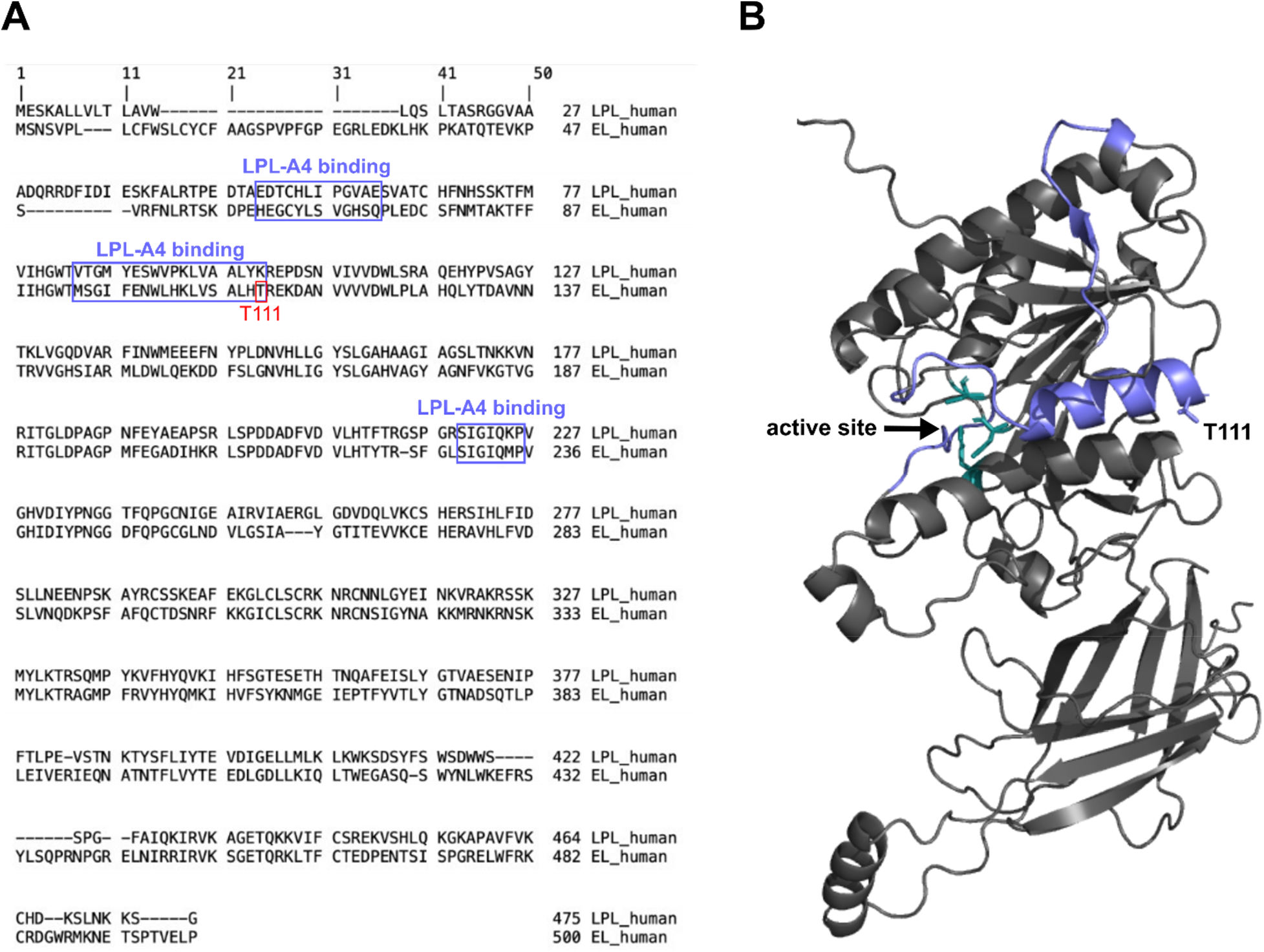
Potential ANGPTL binding sites on EL. **(A)** MAFFT multiple sequence alignment of human LPL and human EL. ANGPTL4 binding sites on LPL and the homologous regions of EL are boxed in purple. The T111 residue is boxed in red. **(B)** Predicted structure of WT EL indicating the catalytic triad (green) and the regions homologous to the ANGPTL4 binding sites on LPL (purple).

**Figure 4.**
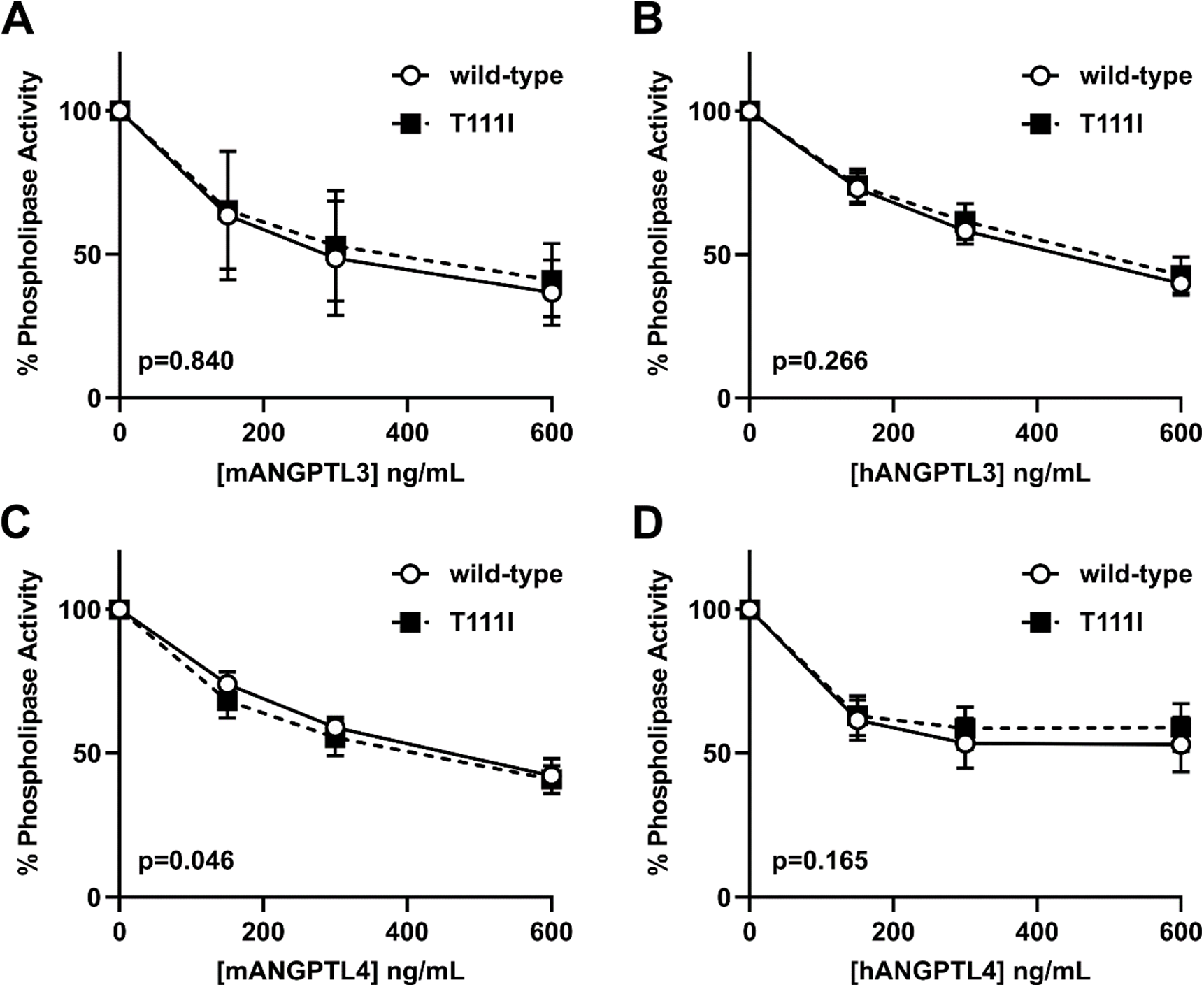
Inhibition of EL T111I by ANGPTL3 and ANGPTL4. WT EL and T111I EL were incubated at 37°C for 30 minutes with increasing concentrations of **(A)** mouse ANGPTL3, **(B)** human ANGPTL3, **(C)** mouse ANGPTL4, or **(D)** human ANGPTL4. Activity was measured using a fluorescence-based phospholipase activity assay. Points represent mean (±SD) of 4 experiments each run with 2 biological duplicates. P values determined by one phase decay least squares fit.

*In vivo*, EL can be bound to the surface of endothelial cells through interactions with heparan sulfate proteoglycans (HSPGs) (55), and we have previously found that EL bound to endothelial cells is resistant to ANGPTL3 inhibition (34). We next therefore asked if the T111I substitution affected the ability of endothelial cell-bound EL to be inhibited by ANGPTL3 or ANGPTL4. We treated rat heart microvessel endothelial cells (RHMVECs) stably expressing WT EL or T111I EL **(Fig. 5A)** with ANGPTL3 or ANGPTL4 and measured lipase activity. As expected, EL bound to the endothelial cell surface was resistant to ANGPTL3 and ANGPTL4 inhibition **(Fig 5)**. However, we again found little difference in the ability of ANGPTL3 **(Fig. 5B-C)** or ANGPTL4 **(Fig. 5D-E)** to inhibit T111I EL compared with wild-type EL.

**Figure 5.**
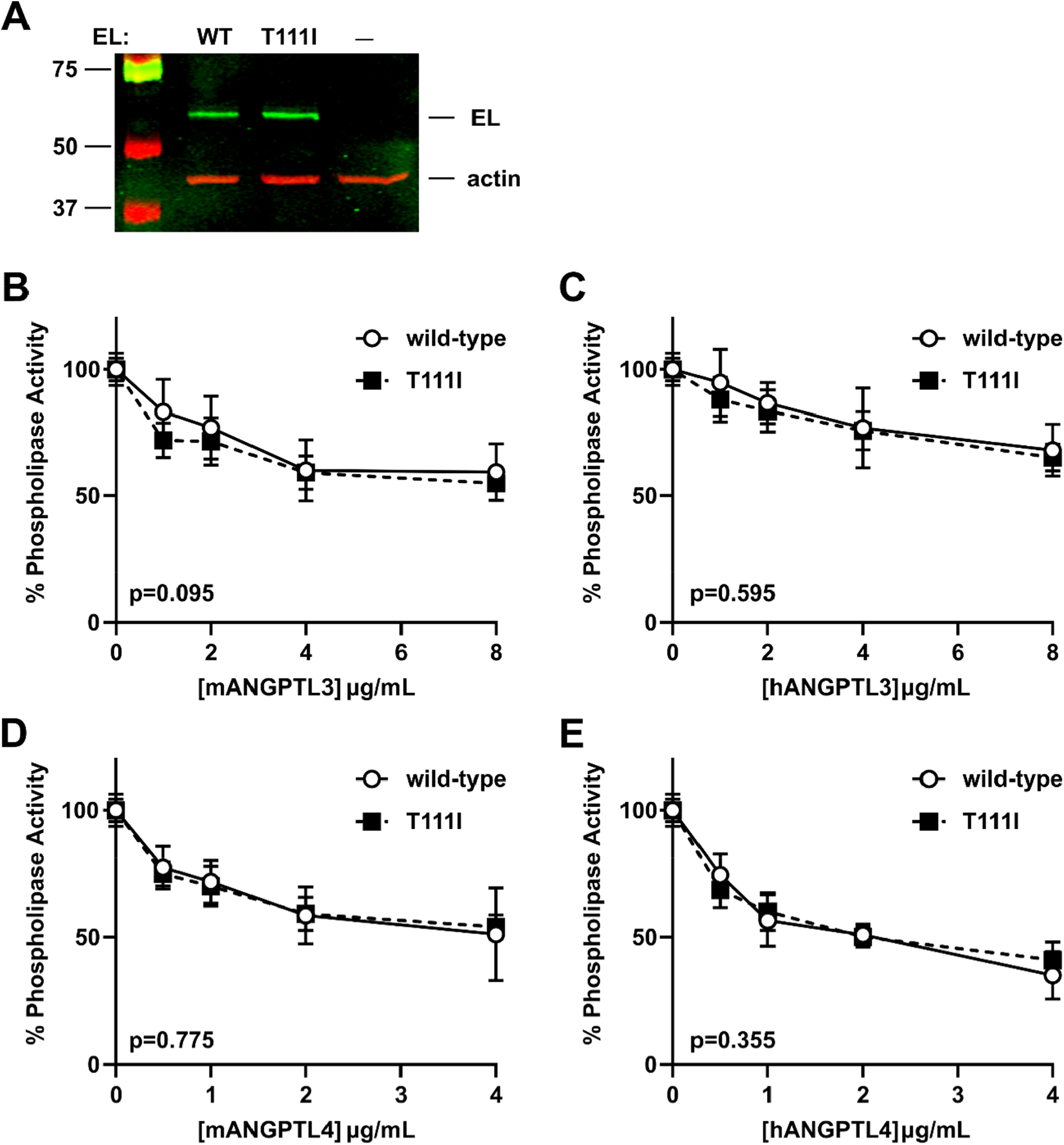
Inhibition of membrane-tethered EL T111I by ANGPTL3 and ANGPTL4. **(A)** Expression of WT and T111I EL from transduced RHMVEC. Western blot shows expression of EL in the lysate of virally transduced endothelial cells. **(B-E)** WT EL and T111I EL expressing RHMVEC were incubated at 37°C for 30 minutes with increasing concentrations of (**B**) mouse ANGPTL3, **(C)** human ANGPTL3, **(D)** mouse ANGPTL4, or **(E)** human ANGPTL4. Activity was measured using a fluorescence-based phospholipase activity assay. Points represent mean (±SD) of 4 experiments each run with 2 biological duplicates. P values determined by one phase decay least squares fit.

## DISCUSSION

In this study, we evaluated a common EL variant, T111I, for its effects on EL activity, stability, and ability to be inhibited by ANGPTL proteins. In all these parameters, we found that EL T111I was no different than wildtype EL.

The 584C/T polymorphism, which encodes the T111I mutation in EL, has been associated with increased HDL levels in some population studies. The variant was originally discovered by sequencing black and Caucasian individuals with high HDL-C levels (11). Subsequently, several groups found an association between the T111I variant and HDL-C levels and other cardiovascular risk factors (10, 12–17), but several other studies found no association (18–23). Numerous studies continue to be performed to evaluate a potential association between this EL variant, HDL-C levels, and the risk of CAD. A meta-analysis, performed by Cai *et al*. in 2014, included 9 case-control studies of Caucasian and Asian populations. This meta-analysis, which considered the sample size of the previously published studies and repeated the statistical analysis, found that carriers of the 584C/T polymorphism did have higher HDL-C levels compared to non-carriers but did not find a significant association between the variant and the risk of coronary heart disease (17). A second meta-analysis performed by Zhao *et al*. in 2020 included 13 studies from countries, including The United States, China, Japan, Netherlands, Iran, Egypt, and Turkey. This analysis did not examine association with HDL-C levels, but did find that EL 584C/T was associated with CAD susceptibility (56). A third meta-analysis by Wu *et al*. included 14 case-control studies with 9731 subjects (57). This meta-analysis also found a significant association between the EL 584C/T polymorphism and CAD, but only in African populations (57). Again HDL-C levels were not analyzed. In summary, the association between the EL T111I variant and HDL-C or the risk of CAD remains somewhat unsettled.

If T111I is indeed associated with high HDL levels, the simplest explanation is that it alters EL function. Functional changes could include changes in enzymatic activity, changes in stability, and changes in interactions with other proteins, such as the endogenous inhibitor ANGPTL3. A previous study by Edmonson *et al*. found no difference in phospholipase activity between wild-type and T111I EL when tested either by measuring the hydrolysis of dipalmitoylphosphatidyl choline or by using isolated human HDL_3_ as a substrate (23). We likewise found no differences in phospholipase activity using either a fluorescence-based assay or fatty-acid release from human HDL particles.

Although the structure of EL has not been solved experimentally, the ColabFold structure puts the T111 at the end of an alpha helix in the N-terminal domain of EL. Although the N-terminal domain contains the catalytic domain, T111 is not near the active site, nor does it seem positioned to contribute to proper folding of EL. Using both computational and experimental techniques, we found no evidence that the T111I mutation disrupts EL stability, at least in vitro. These data support previous predictions which utilized homology modelling to predict the structure of EL and CHARMM calculation protocols to predict stability changes and suggested that the T111I mutation does not have any significant effects on the EL protein stability (40).

The location of T111 on the outer face of an alpha helix on EL suggests that it could potentially be involved in interactions with other proteins. Indeed, in the homologous lipase, LPL, this same alpha helix is thought to interact with the lipase inhibitor ANGPTL4 (42, 52). However, it is important to note that the T111 is not conserved between EL and LPL. Moreover, we did not observe any meaningful differences in inhibition by either ANGPTL3 or ANGPTL4 when threonine 111 was mutated to isoleucine. We conclude that if T111 is involved in protein binding, it is not with ANGPTL proteins.

The data in this paper suggest that the T111I variation does not alter EL specific activity, protein stability, or inhibition by ANGPTL proteins. However, it is important to note that it is also possible that T111I induces small enough changes in EL activity, stability, or inhibition by ANGPTL proteins that the sensitivity and reproducibility of our assays were unable to detect these differences. Our findings do not rule out the possibility that T111I compromises EL function. It is possible that T111I alters the interactions of EL with proteins other than ANGPTLs. The T111I variant introduces a large hydrophobic residue on the surface of EL, which would increase the propensity for non-specific hydrophobic associations, and these interactions could potentially alter EL function. The T111I variant could also alter posttranslational modifications or alter the behavior of EL *in vivo* in ways that cannot be easily tested in *in vitro*. Alternatively, the T111I allele may be associated with increased cardiovascular risk, but not causal, with causality coming from other alleles in the same haplotype (22, 40).

## Abbreviations

ANGPTL: angiopoietin-like
CAD: coronary artery disease
CM: control media
EL: endothelial lipase
GPIHBP1: glycosylphosphatidylinositol-anchored HDL binding protein 1
HEK: human embryonic kidney
HSPG: heparan-sulfate proteoglycan
PEI: polyethylenimine
RHMVECs: rat heart microvessel endothelial cells

## REFERENCES

1. Tall, A. R. 1998. An overview of reverse cholesterol transport. Eur Heart J. 19 Suppl A: A31-35.

2. Wang, N., D. Lan, W. Chen, F. Matsuura, and A. R. Tall. 2004. ATP-binding cassette transporters G1 and G4 mediate cellular cholesterol efflux to high-density lipoproteins. Proc Natl Acad Sci U S A. 101: 9774–9779.

3. Larrede, S., C. M. Quinn, W. Jessup, E. Frisdal, M. Olivier, V. Hsieh, M.-J. Kim, M. Van Eck, Couvert, A. Carrie, P. Giral, M. J. Chapman, M. Guerin, and W. Le Goff. 2009. Stimulation of cholesterol efflux by LXR agonists in cholesterol-loaded human macrophages is ABCA1-dependent but ABCG1-independent. Arterioscler Thromb Vasc Biol. 29: 1930–1936.

4. Tarling, E. J., and P. A. Edwards. 2011. ATP binding cassette transporter G1 (ABCG1) is an intracellular sterol transporter. Proc Natl Acad Sci U S A. 108: 19719–19724.

5. Bowry, V. W., K. K. Stanley, and R. Stocker. 1992. High density lipoprotein is the major carrier of lipid hydroperoxides in human blood plasma from fasting donors. Proc Natl Acad Sci U S A. 89: 10316–10320.

6. Hirata, K., H. L. Dichek, J. A. Cioffi, S. Y. Choi, N. J. Leeper, L. Quintana, G. S. Kronmal, A. D. Cooper, and T. Quertermous. 1999. Cloning of a unique lipase from endothelial cells extends the lipase gene family. J Biol Chem. 274: 14170–14175.

7. Jahangiri, A., D. J. Rader, D. Marchadier, L. K. Curtiss, D. J. Bonnet, and K.-A. Rye. 2005. Evidence that endothelial lipase remodels high density lipoproteins without mediating the dissociation of apolipoprotein A-I. J Lipid Res. 46: 896–903.

8. Jaye, M., K. J. Lynch, J. Krawiec, D. Marchadier, C. Maugeais, K. Doan, V. South, D. Amin, M. Perrone, and D. J. Rader. 1999. A novel endothelial-derived lipase that modulates HDL metabolism. Nature genetics. 21: 424–428.

9. Ishida, T., S. Choi, R. K. Kundu, K. Hirata, E. M. Rubin, A. D. Cooper, and T. Quertermous. 2003. Endothelial lipase is a major determinant of HDL level. J Clin Invest. 111: 347–355.

10. Ma, K., M. Cilingiroglu, J. D. Otvos, C. M. Ballantyne, A. J. Marian, and L. Chan. 2003. Endothelial lipase is a major genetic determinant for high-density lipoprotein concentration, structure, and metabolism. Proceedings of the National Academy of Sciences. 100: 2748–2753.

11. deLemos, A. S., M. L. Wolfe, C. J. Long, R. Sivapackianathan, and D. J. Rader. 2002. Identification of genetic variants in endothelial lipase in persons with elevated high-density lipoprotein cholesterol. Circulation. 106: 1321–1326.

12. Mank-Seymour, A. R., K. L. Durham, J. F. Thompson, A. B. Seymour, and P. M. Milos. 2004. Association between single-nucleotide polymorphisms in the endothelial lipase (LIPG) gene and high-density lipoprotein cholesterol levels. Biochim Biophys Acta. 1636: 40–46.

13. Hutter, C. M., M. A. Austin, F. M. Farin, H.-M. Viernes, K. L. Edwards, D. L. Leonetti, M. J. McNeely, and W. Y. Fujimoto. 2006. Association of endothelial lipase gene (LIPG) haplotypes with high-density lipoprotein cholesterol subfractions and apolipoprotein AI plasma levels in Japanese Americans. Atherosclerosis. 185: 78–86.

14. Tang, N.-P., L.-S. Wang, L. Yang, B. Zhou, H.-J. Gu, Q.-M. Sun, R.-H. Cong, H.-J. Zhu, and B. Wang. 2008. Protective effect of an endothelial lipase gene variant on coronary artery disease in a Chinese population. J Lipid Res. 49: 369–375.

15. Paradis, M.-E., P. Couture, Y. Bosse, J.-P. Despres, L. Perusse, C. Bouchard, M.-C. Vohl, and B. Lamarche. 2003. The T111I mutation in the EL gene modulates the impact of dietary fat on the HDL profile in women. J Lipid Res. 44: 1902–1908.

16. Solim, L. A., I. A. Gencan, B. Çelik, A. Ataacar, U. Koç, B. Büyükören, G. Güngör, and S. Isbir. 2018. Endothelial Lipase Gene Polymorphism (584 C/T) in Coronary Artery Patients Among a Turkish Population. In Vivo. 32: 1105–1109.

17. Cai, G., Z. Huang, B. Zhang, W. Weng, and G. Shi. 2014. The associations between endothelial lipase 584C/T polymorphism and HDL-C level and coronary heart disease susceptibility: a meta-analysis. Lipids Health Dis. 13: 85.

18. Halverstadt, A., D. A. Phares, R. E. Ferrell, K. R. Wilund, A. P. Goldberg, and J. M. Hagberg. 2003. High-density lipoprotein-cholesterol, its subfractions, and responses to exercise training are dependent on endothelial lipase genotype. Metabolism. 52: 1505–1511.

19. Shimizu, M., K. Kanazawa, K. Hirata, T. Ishida, E. Hiraoka, Y. Matsuda, C. Iwai, Y. Miyamoto, M. Hashimoto, T. Kajiya, H. Akita, and M. Yokoyama. 2007. Endothelial lipase gene polymorphism is associated with acute myocardial infarction, independently of high-density lipoprotein-cholesterol levels. Circ J. 71: 842–846.

20. Jensen, M. K., E. B. Rimm, K. J. Mukamal, A. C. Edmondson, D. J. Rader, U. Vogel, A. Tjønneland, T. I. A. Sørensen, E. B. Schmidt, and K. Overvad. 2009. The T111I variant in the endothelial lipase gene and risk of coronary heart disease in three independent populations. Eur Heart J. 30: 1584–1589.

21. Yamakawa-Kobayashi, K., H. Yanagi, K. Endo, T. Arinami, and H. Hamaguchi. 2003. Relationship between serum HDL-C levels and common genetic variants of the endothelial lipase gene in Japanese school-aged children. Hum Genet. 113: 311–315.

22. Razzaghi, H., S. A. Santorico, and M. I. Kamboh. 2012. Population-Based Resequencing of LIPG and ZNF202 Genes in Subjects with Extreme HDL Levels. Front Genet. 3: 89.

23. Edmondson, A. C., R. J. Brown, S. Kathiresan, L. A. Cupples, S. Demissie, A. K. Manning, M. K. Jensen, E. B. Rimm, J. Wang, A. Rodrigues, V. Bamba, S. A. Khetarpal, M. L. Wolfe, S. DerOhannessian, M. Li, M. P. Reilly, J. Aberle, D. Evans, R. A. Hegele, and D. J. Rader. 2009. Loss-of-function variants in endothelial lipase are a cause of elevated HDL cholesterol in humans. J Clin Invest. 119: 1042–1050.

24. Smith, C. E., D. K. Arnett, M. Y. Tsai, C.-Q. Lai, L. D. Parnell, J. Shen, M. Laclaustra, M. Junyent, and J. M. Ordovás. 2009. Physical inactivity interacts with an endothelial lipase polymorphism to modulate high density lipoprotein cholesterol in the GOLDN study. Atherosclerosis. 206: 500–504.

25. Shimamura, M., M. Matsuda, H. Yasumo, M. Okazaki, K. Fujimoto, K. Kono, T. Shimizugawa, Y. Ando, R. Koishi, T. Kohama, and others. 2007. Angiopoietin-like protein3 regulates plasma HDL cholesterol through suppression of endothelial lipase. Arteriosclerosis, thrombosis, and vascular biology. 27: 366–372.

26. Chi, X., E. C. Britt, H. W. Shows, A. J. Hjelmaas, S. K. Shetty, E. M. Cushing, W. Li, A. Dou, R. Zhang, and B. S. J. Davies. 2017. ANGPTL8 promotes the ability of ANGPTL3 to bind and inhibit lipoprotein lipase. Molecular Metabolism. 6: 1137–1149.

27. Lee, E.-C., U. Desai, G. Gololobov, S. Hong, X. Feng, X.-C. Yu, J. Gay, N. Wilganowski, C. Gao, L.-L. Du, J. Chen, Y. Hu, S. Zhao, L. Kirkpatrick, M. Schneider, B. P. Zambrowicz, G. Landes, D. R. Powell, and W. K. Sonnenburg. 2009. Identification of a New Functional Domain in Angiopoietin-like 3 (ANGPTL3) and Angiopoietin-like 4 (ANGPTL4) Involved in Binding and Inhibition of Lipoprotein Lipase (LPL). J Biol Chem. 284: 13735–13745.

28. Shimizugawa, T., M. Ono, M. Shimamura, K. Yoshida, Y. Ando, R. Koishi, K. Ueda, T. Inaba, H. Minekura, T. Kohama, and H. Furukawa. 2002. ANGPTL3 Decreases Very Low Density Lipoprotein Triglyceride Clearance by Inhibition of Lipoprotein Lipase. J. Biol. Chem. 277: 33742–33748.

29. Shan, L., X.-C. Yu, Z. Liu, Y. Hu, L. T. Sturgis, M. L. Miranda, and Q. Liu. 2009. The angiopoietin-like proteins ANGPTL3 and ANGPTL4 inhibit lipoprotein lipase activity through distinct mechanisms. J. Biol. Chem. 284: 1419–1424.

30. Sonnenburg, W. K., D. Yu, E.-C. Lee, W. Xiong, G. Gololobov, B. Key, J. Gay, N. Wilganowski, Y. Hu, S. Zhao, M. Schneider, Z.-M. Ding, B. P. Zambrowicz, G. Landes, D. R. Powell, and U. Desai. 2009. GPIHBP1 stabilizes lipoprotein lipase and prevents its inhibition by angiopoietin-like 3 and angiopoietin-like 4. J Lipid Res. 50: 2421–2429.

31. Yau, M., Y. Wang, K. S. L. Lam, J. Zhang, D. Wu, and A. Xu. 2009. A Highly Conserved Motif within the NH2-terminal Coiled-coil Domain of Angiopoietin-like Protein 4 Confers Its Inhibitory Effects on Lipoprotein Lipase by Disrupting the Enzyme Dimerization. J Biol Chem. 284: 11942–11952.

32. Jin, W., X. Wang, J. S. Millar, T. Quertermous, G. H. Rothblat, J. M. Glick, and D. J. Rader. 2007. Hepatic proprotein convertases modulate HDL metabolism. Cell Metab. 6: 129–136.

33. Chi, X., S. K. Shetty, H. W. Shows, A. J. Hjelmaas, E. K. Malcolm, and B. S. J. Davies. 2015. Angiopoietin-like 4 Modifies the Interactions between Lipoprotein Lipase and Its Endothelial Cell Transporter GPIHBP1. J. Biol. Chem. 290: 11865–11877.

34. Sylvers-Davie, K. L., A. Segura-Roman, A. M. Salvi, K. J. Schache, and B. S. J. Davies. 2021. Angiopoietin-like 3 Inhibition of Endothelial Lipase Is Not Modulated by Angiopoietin-like 8. J Lipid Res. 100112.

35. Dull, T., R. Zufferey, M. Kelly, R. J. Mandel, M. Nguyen, D. Trono, and L. Naldini. 1998. A third-generation lentivirus vector with a conditional packaging system. J Virol. 72: 8463–8471.

36. Basu, D., X. Lei, J. Josekutty, M. M. Hussain, and W. Jin. 2013. Measurement of the phospholipase activity of endothelial lipase in mouse plasma. J Lipid Res. 54: 282–289.

37. Mirdita, M., K. Schütze, Y. Moriwaki, L. Heo, S. Ovchinnikov, and M. Steinegger. 2021. ColabFold - Making protein folding accessible to all. [online] https://www.biorxiv.org/content/10.1101/2021.08.15.456425v2 (Accessed November 26, 2021).

38. Jumper, J., R. Evans, A. Pritzel, T. Green, M. Figurnov, O. Ronneberger, K. Tunyasuvunakool, R. Bates, A. Žídek, A. Potapenko, A. Bridgland, C. Meyer, S. A. A. Kohl, A. J. Ballard, A. Cowie, B. Romera-Paredes, S. Nikolov, R. Jain, J. Adler, T. Back, S. Petersen, D. Reiman, E. Clancy, M. Zielinski, M. Steinegger, M. Pacholska, T. Berghammer, S. Bodenstein, D. Silver, O. Vinyals, A. W. Senior, K. Kavukcuoglu, P. Kohli, and D. Hassabis. 2021. Highly accurate protein structure prediction with AlphaFold. Nature. 596: 583–589.

39. UniProt Consortium. 2021. UniProt: the universal protein knowledgebase in 2021. Nucleic Acids Res. 49: D480–D489.

40. Razzaghi, H., A. Tempczyk-Russell, K. Haubold, S. A. Santorico, T. Shokati, U. Christians, and M. E. A. Churchill. 2013. Genetic and structure-function studies of missense mutations in human endothelial lipase. PLoS ONE. 8: e55716.

41. Rodrigues, C. H. M., D. E. V. Pires, and D. B. Ascher. 2021. DynaMut2: Assessing changes in stability and flexibility upon single and multiple point missense mutations. Protein Sci. 30: 60–69.

42. Leth-Espensen, K. Z., K. K. Kristensen, A. Kumari, A.-M. L. Winther, S. G. Young, T. J. D. Jørgensen, and M. Ploug. 2021. The intrinsic instability of the hydrolase domain of lipoprotein lipase facilitates its inactivation by ANGPTL4-catalyzed unfolding. Proc Natl Acad Sci U S A. 118: e2026650118.

43. Mysling, S., K. K. Kristensen, M. Larsson, O. Kovrov, A. Bensadouen, T. J. Jørgensen, G. Olivecrona, S. G. Young, and M. Ploug. The angiopoietin-like protein ANGPTL4 catalyzes unfolding of the hydrolase domain in lipoprotein lipase and the endothelial membrane protein GPIHBP1 counteracts this unfolding. eLife. 5. [online] https://www.ncbi.nlm.nih.gov/pmc/articles/PMC5148603/.

44. Ge, H., G. Yang, X. Yu, T. Pourbahrami, and C. Li. 2004. Oligomerization state-dependent hyperlipidemic effect of angiopoietin-like protein 4. J Lipid Res. 45: 2071–2079.

45. Lichtenstein, L., F. Mattijssen, N. J. de Wit, A. Georgiadi, G. J. Hooiveld, R. van der Meer, Y. He, L. Qi, A. Köster, J. T. Tamsma, N. S. Tan, M. Müller, and S. Kersten. 2010. Angptl4 protects against severe proinflammatory effects of saturated fat by inhibiting fatty acid uptake into mesenteric lymph node macrophages. Cell Metab. 12: 580–592.

46. Mandard, S., F. Zandbergen, E. van Straten, W. Wahli, F. Kuipers, M. Müller, and S. Kersten. 2006. The Fasting-induced Adipose Factor/Angiopoietin-like Protein 4 Is Physically Associated with Lipoproteins and Governs Plasma Lipid Levels and Adiposity. J. Biol. Chem. 281: 934–944.

47. Köster, A., Y. B. Chao, M. Mosior, A. Ford, P. A. Gonzalez-DeWhitt, J. E. Hale, D. Li, Y. Qiu, C. C. Fraser, D. D. Yang, J. G. Heuer, S. R. Jaskunas, and P. Eacho. 2005. Transgenic angiopoietin-like (angptl)4 overexpression and targeted disruption of angptl4 and angptl3: regulation of triglyceride metabolism. Endocrinology. 146: 4943–4950.

48. Sukonina, V., A. Lookene, T. Olivecrona, and G. Olivecrona. 2006. Angiopoietin-like protein 4 converts lipoprotein lipase to inactive monomers and modulates lipase activity in adipose tissue. Proc. Natl. Acad. Sci. U.S.A. 103: 17450–17455.

49. Haller, J. F., I. J. Mintah, L. M. Shihanian, P. Stevis, D. Buckler, C. A. Alexa-Braun, S. Kleiner, S. Banfi, J. C. Cohen, H. H. Hobbs, G. D. Yancopoulos, A. J. Murphy, V. Gusarova, and J. Gromada. 2017. ANGPTL8 requires ANGPTL3 to inhibit lipoprotein lipase and plasma triglyceride clearance. J Lipid Res. 58: 1166–1173.

50. Kovrov, O., K. K. Kristensen, E. Larsson, M. Ploug, and G. Olivecrona. 2019. On the mechanism of angiopoietin-like protein 8 for control of lipoprotein lipase activity. Journal of Lipid Research. 60: 783.

51. Chen, Y. Q., T. G. Pottanat, R. W. Siegel, M. Ehsani, Y.-W. Qian, E. Y. Zhen, A. Regmi, W. C. Roell, H. Guo, M. J. Luo, R. E. Gimeno, F. van’t Hooft, and R. J. Konrad. 2020. Angiopoietin-like protein 8 differentially regulates ANGPTL3 and ANGPTL4 during postprandial partitioning of fatty acids. Journal of Lipid Research. 61: 1203.

52. Kristensen, K. K., K. Z. Leth-Espensen, H. D. T. Mertens, G. Birrane, M. Meiyappan, G. Olivecrona, T. J. D. Jørgensen, S. G. Young, and M. Ploug. 2020. Unfolding of monomeric lipoprotein lipase by ANGPTL4: Insight into the regulation of plasma triglyceride metabolism. Proc Natl Acad Sci U S A. 117: 4337–4346.

53. Katoh, K., and D. M. Standley. 2013. MAFFT multiple sequence alignment software version 7: improvements in performance and usability. Mol Biol Evol. 30: 772–780.

54. Chen, Y. Q., T. G. Pottanat, R. W. Siegel, M. Ehsani, Y.-W. Qian, and R. J. Konrad. 2021. Angiopoietin-like protein 4 (ANGPTL4) is an inhibitor of endothelial lipase (EL) while the ANGPTL4/8 complex has reduced EL-inhibitory activity. Heliyon. 7: e07898.

55. Fuki, I. V., N. Blanchard, W. Jin, D. H. L. Marchadier, J. S. Millar, J. M. Glick, and D. J. Rader. 2003. Endogenously produced endothelial lipase enhances binding and cellular processing of plasma lipoproteins via heparan sulfate proteoglycan-mediated pathway. J. Biol. Chem. 278: 34331–34338.

56. Zhao, H., R. Zhao, S. Hu, and J. Rong. 2020. Gene polymorphism associated with angiotensinogen (M235T), endothelial lipase (584C/T) and susceptibility to coronary artery disease: a meta-analysis. Biosci Rep. 40: BSR20201414.

57. Wu, Y., L. Ma, H. Zhang, X. Chen, X. Xu, and Z. Hu. 2020. Significant association between the endothelial lipase gene 584C/T polymorphism and coronary artery disease risk. Biosci Rep. 40: BSR20200027.

